# Hidden introductions of freshwater red algae via the aquarium trade exposed by DNA barcodes

**DOI:** 10.1101/2020.06.30.180042

**Authors:** Shing Hei Zhan, Tsai-Yin Hsieh, Lan-Wei Yeh, Ting-Chun Kuo, Shoichiro Suda, Shao-Lun Liu

## Abstract

The global aquarium trade can introduce alien freshwater invaders, potentially impacting local aquatic ecosystems and their biodiversity. The role of the aquarium trade in spreading freshwater red macroalgae that hitchhike on ornamental aquatic plants and animals is unassessed. We investigated this human-mediated phenomenon via a broad biodiversity survey and genetic analysis of freshwater red algae in the field and aquarium shops in East Asia. Using *rbc*L-based DNA barcoding, we surveyed 125 samples from 46 field sites and 88 samples from 53 aquarium shops (213 samples in total) mostly across Taiwan – a key hub in the global aquarium trade – as well as in Hong Kong, Okinawa (Japan), the Philippines, and Thailand. We augmented our *rbc*L sequences with GenBank *rbc*L sequences that represent 40 additional countries globally. We found 26 molecular operational taxonomic units (mOTUs) in Taiwan, some of which are cryptic. Phylogeographical analysis revealed three potential introduced mOTUs in Taiwan, which exhibit no local genetic variation in Taiwan and are distributed across continents. Also, we posit that some presumably endangered freshwater red algae may be preserved in aquaria, an unintentional *ex situ* conservation site for these organisms that are vulnerable to water pollution from anthropogenic disturbances. Collectively, these data suggest that freshwater red algae have been hitchhiking and dispersed via the aquarium trade, an important and overlooked mechanism of introduction of these organisms across the globe.

## INTRODUCTION

The aquarium trade facilitates introduction of alien species and homogenization of biodiversity in aquatic ecosystems worldwide (e.g., Padilla & William, 2004; Rahel, 2007; Strechker et al., 2011). Many introduced species have hardly discernible or subtle impacts on the biodiversity of local environments, but some might become harmful invasive species (Strayer, 2010; Simberloff, 2014). Efforts to eradicate invasive species incur substantial social and economic costs, yet they are often unsuccessful (reviewed in Simberloff et al., 2013). Therefore, to prevent and control the spread of invasive species, it is important to establish an early warning and rapid response system. To support the implementation of such a system, knowledge about the taxonomic diversity and transportation of potential invasive species is critically needed.

As invasive species, aquatic hitchhikers are damaging to marine and freshwater ecosystems and potentially to public health (e.g., Patoka et al., 2016; Duggan & Pullan, 2017; Duggan et al., 2018). It has been suggested that the socioeconomic impact of aquatic hitchhikers, such as algae, is often overlooked (reviewed in Kaštovský et al., 2010). For instance, an introduced green alga, *Hydrodictyon reticulatum*, has been reported to cause the decline of local fish and aquatic plants through habitat transformation, but it had been unnoticed for long until its blooming (Wells et al., 1999; Wells & Clayton, 2001).

Unlike ornamental animals and plants in the aquarium trade, less attention has been paid to the diversity and introduction potential of aquatic hitchhikers largely because of their species crypticity (Stoyneva et al, 2006; Kato et al., 2009). Freshwater red macroalgae are aquatic hitchhikers that are occasionally found in aquarium tanks (e.g., Kaufmann, 2010), but these algae are rare in the field near cities due to their vulnerability to polluted water (Sheath & Hambrook, 1990; Sheath & Vis, 2015). These algae produce inconspicuous spores that suspend in water or adhere to aquatic organisms, such as aquatic plants (**Fig. 1a**), crayfish (**Fig. 1b**), and snails (**Fig. 1c-e**). Spores can be generated by asexual reproduction (when the algae are turf-like sporophytes, or chantransia) or by sexual reproduction (when the algae appear as thread-like, mucilaginous gametophytes). The asexual form may facilitate the population establishment of introduced freshwater red algae in the non-native range where mating partners may be unavailable, but to our knowledge this has not been supported empirically.

**Figure 1.**
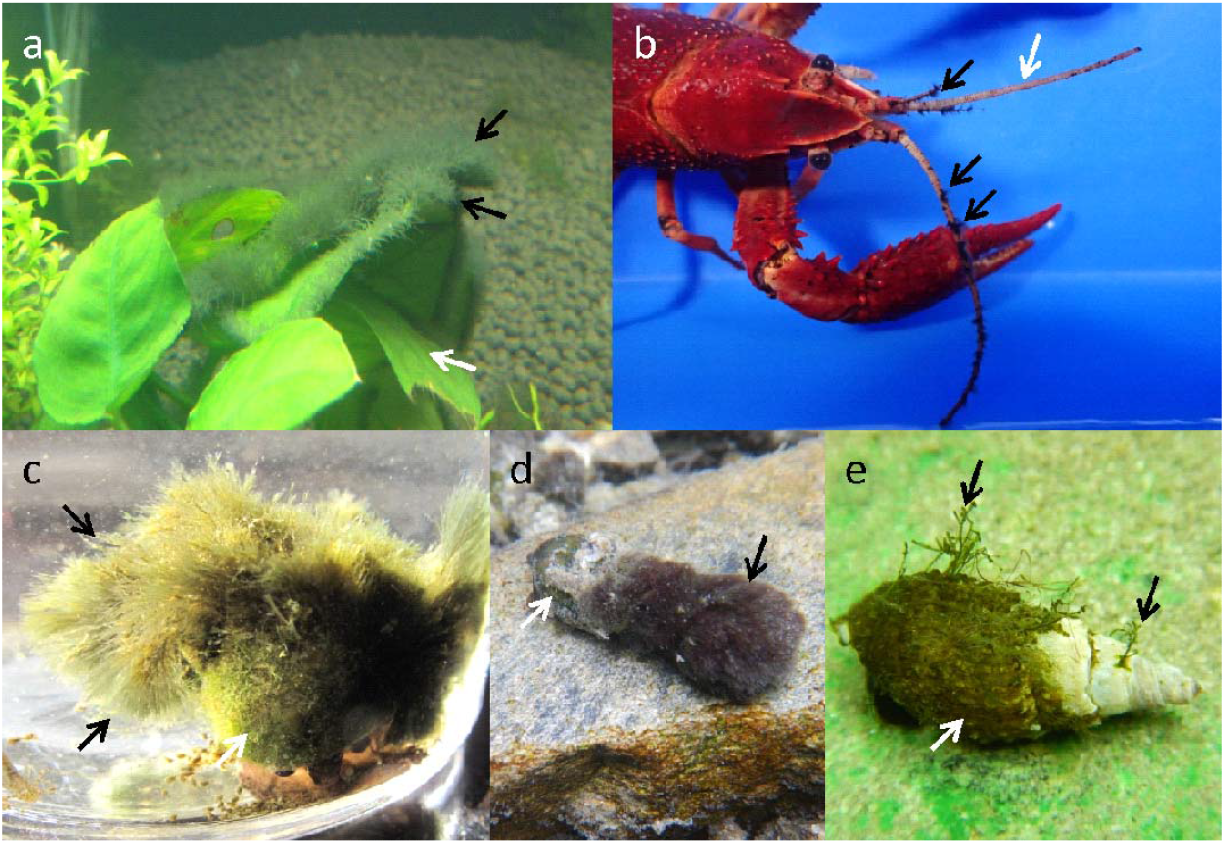
Examples of epiphytic (a) and epizoic (b-e) freshwater red macroalgae as aquatic hitchhikers (black arrows). (a) *Compsopogon coeruleus* (THU.369) (black arrow) on an ornamental aquatic plant, *Aubulias barteri* (white arrow), in the Gou-Lao-Ban Pet Shop (Taoyuan, Taiwan). (b) The chantransia stage of *Sheathia dispersa* (THU.575) (black arrow) on a red swamp crayfish, *Procambarus clarkii* (white arrow), in the Yu-Zhong-Yu Aquarium Shop (Taichung, Taiwan). (c) The chantransia stage of *Montagnia macrospora* (THU.537) (black arrow) on an apple snail, *Pomacea canaliculata* (white arrow), in Nanshi River (Taichung, Taiwan). (d) The chantransia stage of *Montagnia macrospora* (THU.401) (black arrow) on a chopstick snail, *Stenomelania* sp. (white arrow), in the Mataian Wetland Ecological Park (Hualien, Taiwan). (e) A six-month lab culture showing the growth of the gametophyte of *S. dispersa* (black arrow) on the shell of a chopstick snail, *Stenomelania* sp. (white arrow), which was collected from the field.

Only a few freshwater algal species have been documented as biological invasions. These known invaders are *Bangia atropurpurea* (a red alga; Lin & Blum, 1977), *Chara connivens* (a green alga; Luther, 1979), *Nitellopsis obtusa* (a green alga; Schloesser, 1986), *Compsopogon caeruleus* (a red alga; Manny et al., 1991), *Hydrodictyon reticulatum* (a green alga; Hawes et al., 1991), and *Ulva flexuosa* (a green alga; Kaštovský et al., 2010). These algae have been introduced by discharge of ballast water (e.g., Lin & Blum, 1977; Manny et al., 1991) or accidental release from laboratory experiments (Hawes et al., 1991). Freshwater red macroalgae can be dispersed through the aquarium trade (*Composopogon caeruleus*, Stoyneva et al., 2006; *Montagnia macrospora*, Kato et al., 2009). Recently, it has been speculated that the cosmopolitan distribution of freshwater red macroalgae may be facilitated by the aquarium trade (e.g., Carlile & Sherwood, 2013; Johnston et al., 2018). Beyond speculation, it has not been explored whether the aquarium trade can serve as a vector for the global dispersal of freshwater red algae.

Traditional biodiversity monitoring programs depend on morphology-based approaches, which require time-consuming diagnostics by taxonomy experts (Riedel et al., 2013). Hitchhiking freshwater red macroalgae are morphologically indistinguishable, thus rendering their species identification challenging. A powerful alternative approach to monitor biodiversity is DNA barcoding, which typically involves sequencing individual genetic loci, such as *rbc*L. DNA barcodes have been applied to detect introduced and invasive organisms (Armstrong & Ball, 2005; Pečnikar & Buzan, 2014). Recently, Vranken and coworkers (2018) investigated the spread of introduced and invasive marine macroalgae (i.e., seaweeds) via the aquarium trade in Europe using DNA barcodes. No study has examined the introduction of freshwater red macroalgae via the aquarium trade at a geographically broad scale.

DNA barcode sequence data can be utilized to estimate the effect of introduction(s) on the genetic diversity of an alien species (e.g., Bonett et al., 2007; Kinziger et al., 2011). The population of a newly introduced species is predicted to harbor lower genetic diversity than its source population(s) as a consequence of a recent genetic bottleneck (Nei et al., 1975; Barrett & Husband, 1990), but multiple introductions may increase the genetic diversity of an introduced population (Novak & Mack 1993; Novak & Mack 2005) (empirical evidence reviewed in Dlugosch & Parker, 2008). Following this logic, the genetic information in DNA barcodes, in combination with geographic occurrence data, can enable detection of putative introduced or invasive species.

In this study, we employed DNA barcoding to survey the biodiversity of freshwater red macroalgae from the field and aquarium shops across Taiwan – a key hub in the global aquarium trade (Padilla & William, 2004) – as well as in a few surrounding regions (Hong Kong, Okinawa in Japan, the Philippines, and Thailand). We collected 213 macroalgal specimens from 99 sites (mostly in Taiwan, 196 specimens from 90 sites), determined their *rbc*L sequences, and estimated the number and species identity of molecular operational taxonomic units (mOTUs). These data yielded, for the first time, a detailed picture of the biodiversity of freshwater red macroalgae in Taiwan – in the field and in aquaria. To identify potential introduced species in Taiwan, we examined the genetic diversity and global distribution of the mOTUs by analyzing our data jointly with previously published *rbc*L sequences of specimens collected worldwide. Also, we assessed the relative proportion of freshwater red algae in asexual and sexual forms in both the field and aquaria to test whether the asexual form is more frequently observed in aquaria. Finally, we examined ornamental fish trade data to assess the plausibility of an introduction route of an alga via Taiwan’s aquarium trade.

## MATERIALS AND METHODS

### Sample collection

In total, 213 specimens were collected from the field and aquarium shops in Taiwan, Okinawa (Japan), Hong Kong, the Philippines, and Thailand (**Fig. 2**; **Table S1**); of the 213 samples, 125 were collected from 46 field sites and 88 from 53 aquarium shops. One sample that was collected from a brackish area was included in the species delimitation analyses, but it was excluded from other analyses (**Table S1**). After collection, a portion of the material (about 100-200 mg) was preserved in silica gel or 95% ethanol for molecular analyses, whereas the rest of the material was preserved in 10-15% formalin solution for morphological examination.

**Figure 2.**
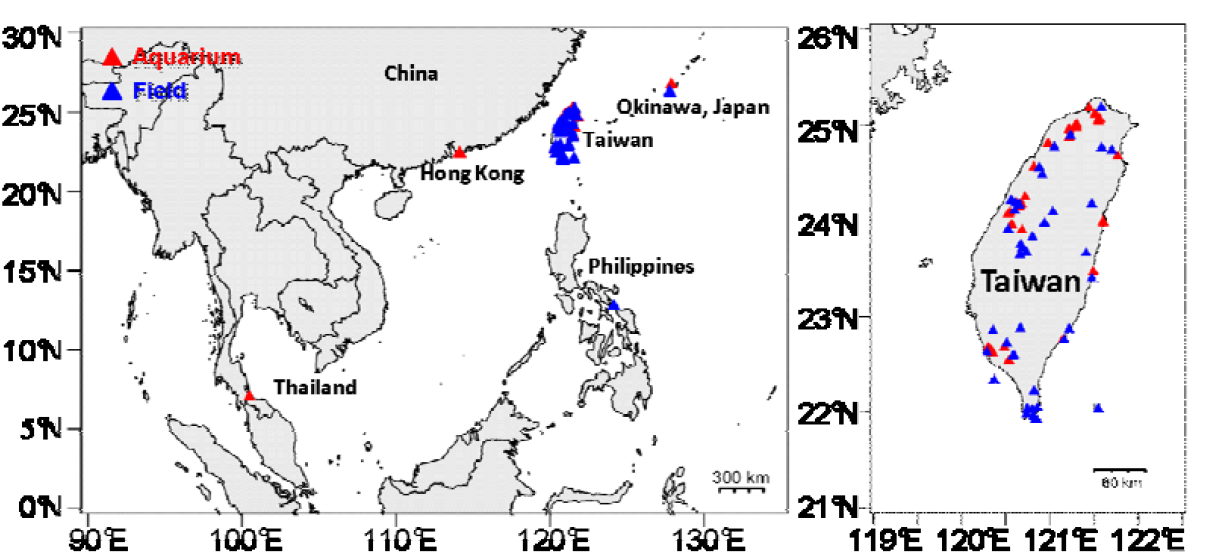
Sites in Taiwan, Okinawa (Japan), Hong Kong, Thailand, and the Philippines sampled in this study.

### DNA extraction, PCR, and Sanger sequencing

DNA was extracted from silica gel-dried specimens or 95% ethanol-preserved specimens by using the commercial ZR Plant/Seed DNA kit (Zymo Research, CA, USA) following the manufacturer’s protocol. We amplified *rbc*L under the PCR conditions described in the protocol of Wang et al. (2005) using these gene-specific primers: two forward (F7: 5’-AACTCTGTAGAACGNACAAG-3’ and F160: 5’-CCTCAACCAGGAGTAGATCC-3’) and one reverse (R753: 5’-GCTCTTTCATACATATCTTCC-3’) (Vis et al., 1998; Lin et al., 2001). The PCR products were sent to the Mission Biotech Company (Taipei, Taiwan) for Sanger sequencing. The GenBank accessions of the newly generated sequences are listed in **Table S1**.

### Sequence acquisition and curation

Poor taxon sampling may lower the accuracy of phylogenetic tree reconstruction and species delimitation (Esselstyn et al., 2012; Ahrens et al., 2016). Therefore, we combined the newly generated *rbc*L sequences with additional *rbc*L sequences from NCBI GenBank (accessed on Oct. 18, 2018). Using the literature as a guide (see the references in **Table S1**), we downloaded from GenBank 1,046 *rbc*L sequences of the freshwater and marine species of the following genera: *Bangia, Bostrychia, Caloglossa*, and *Hildenbrandia.* We built a preliminary phylogeny to identify potential contaminant sequences, which appear as either unexpected non-monophyletic placements or long branch attraction. The following 12 potential contaminant GenBank sequences were removed: ten *Hildenbrandia* sequences (AF107812, AF107818, AF107822, AF107827 to AF107831, AF534406, AF534408), one *Thorea* sequence (GU953248), and one *Bangia* sequence (AF043379).

The 1,034 cleaned GenBank sequences and the 213 newly produced sequences (deposited in GenBank under the accessions: MH835465-MH835677) were combined into a data set of 1,247 sequences (1,054 freshwater taxa and 193 non-freshwater taxa). We reduced sequence redundancy by removing identical sequences using the UCLUST function (hereafter as UCLUST_100_) implemented in USEARCH v8.1.1756 (Edgar, 2010). From each UCLUST_100_ cluster, we randomly selected the longest sequence. This resulted in a final dataset of 639 sequences (500 freshwater taxa, 135 non-freshwater taxa, and 4 taxa that can live in both types of habitat) for the phylogenetic analysis and species delimitation. The sequences were mostly contributed by certain geographical regions – including Australia, Brazil, the United States, Europe, Japan, and Taiwan (primarily from this study) – where there has been extensive taxonomic diversity work (**Fig. S1**). Therefore, the biodiversity reported in this study is biased heavily towards those regions (**Fig. S2**).

We updated species names by consulting the taxonomic nomenclature in AlgaeBase (Guiry & Guiry, 2020; accessed Mar. 7, 2020).

### Phylogenetic tree reconstruction

The *rbc*L sequences were aligned using MUSCLE v3.8.31 with the default settings (Edgar, 2004) followed by manual inspection using BioEdit v7.2.5 (Hall, 1999). Phylogenetic analyses were conducted using two different methods to produce a maximum likelihood (ML) tree and an ultrametric Bayesian tree. A non-ultrametric ML phylogeny was inferred under the best-fit nucleotide substitution model (GTR+G+I; according to the lowest Bayesian Information Criterion) with 1,000 bootstrap replicates using MEGA v6.0 (Tamura et al., 2013). An ultrametric Bayesian phylogeny was estimated using MrBayes v3.2.2 (Ronquist et al., 2012), assuming GTR+G+I (which was determined to be the best-fit model using MEGA) and an Independent Gamma Rate relaxed clock (Lepage et al., 2007). Four MCMC chains (one hot and three cold) were run for 200 million generations, sampling every 20,000 generations and discarding the first 80% of the posterior samples such that the average standard deviation of split frequency was less than 0.01. A majority-rule consensus tree was summarized from the MCMC trees with posterior probabilities as support values using Mesquite v2.75 (Maddison & Maddison, 2011). The ML tree was taken as input for the PTP species delimitation method and the Bayesian consensus tree for the GMYC method (see below).

### Species delimitation

DNA-based algorithmic species delimitation methods are an important tool to differentiate species within morphologically indistinguishable taxonomic groups (reviewed in Leliaert et al., 2014). Several studies have raised concerns about the accuracy of inferring species boundaries using single-locus data (e.g., Knowles & Carstens, 2007; Dupuis et al., 2012). To minimize potential false positives, it is advised to infer mOTUs using multiple species delimitation methods and then to take the most conservative result, which has the fewest mOTUs (Carsten et al., 2013). After determining mOTUs, the non-freshwater taxa were excluded from downstream analyses.

For freshwater red algae, *rbc*L is the only marker with abundantly available sequence data. We estimated mOTUs based on *rbc*L sequences using three different methods: automated barcode gap discovery (ABGD), generalized mixed Yule-coalescent (GMYC), and Poisson tree process (PTP). The distance-based method, ABGD, identifies mOTUs by finding the threshold between intraspecific genetic distances and interspecific genetic distances. The threshold was determined based on the distribution of the pairwise genetic distance between any two given sequences (corrected under the Kimura 2 model). We employed the ABGD online tool (https://bioinfo.mnhn.fr/abi/public/abgd/) using the default settings (Puillandre et al., 2012). Next, we inferred mOTUs using two coalescent-based methods, GMYC and PTP. GMYC determines the point of transition from interspecific branching (a pure birth process) to intraspecific branching (a neutral coalescent process) on a clock-calibrated ultrametric tree (Pons et al., 2006). For the GMYC analysis, the ultrametric tree was imported into Splits (Pons et al., 2006) in R, following the procedure described in Pons et al. (2006). PTP determines mOTUs by detecting the threshold between the intraspecific branching rate and the interspecific branching rate (Zhang et al., 2013). For the PTP analysis, the ML tree was imported into the PTP web server (https://species.h-its.org/).

### Rarefaction analysis

To evaluate sampling effort, we estimated the projected maximum number of mOTUs and sample size-based completeness using iNEXT (<u>iN</u>terpolation and <u>EXT</u>rapolation; Hsieh et al., 2014) in R. The maximum number of mOTUs and the number of samples (i.e., the sample size, or more specifically the number of *rbc*L sequences in this study) needed to detect the maximum number of mOTUs were estimated by a rarefaction analysis. Sample size-based completeness is defined as the number of samples collected in this study divided by the number of samples needed to capture the maximum number of mOTUs.

### Identification of introduced species

Low genetic diversity and large geographical distribution are suggested to be informative criteria to identify potential introduced taxa in the red algae (e.g., Necchi Jr. et al., 2013; Díaz□Tapia et al., 2018; Johnston et al., 2018). We estimated the genetic diversity and geographical range of each mOTU. Two indices of genetic variation, haplotype diversity (Hd) and nucleotide diversity (π), were computed using DnaSP v6 (Rozas et al., 2017). Before calculating these indices, the multiple sequence alignment was trimmed on both ends, where there were overhanging bases (or missing data). Also, we computed the maximum distance between any two field locations for a mOTU as a measure of the geographical range of the mOTU. These data were obtained for the five aquarium mOTUs with sufficient sequence data (at least three aquarium samples and three field samples in Taiwan) to estimate local (i.e., in Taiwan) and global genetic variation. We considered mOTUs with no local genetic variation (Hd = 0 and π = 0) and large geographical range (at least 10,000 km) as likely to be introduced through the aquarium trade in Taiwan.

Additionally, for the introduced mOTUs with enough sampling, their *rbc*L haplotype networks were inferred using PopArt v1.7 (Leigh & Bryant, 2015). Haplotype networks have been utilized to identify the potential geographical source(s) of introductions (e.g., Kato et al., 2009). Populations with higher genetic diversity may be the source of introduced species, which often undergo a genetic bottleneck (e.g., Allendorf & Lundquist, 2003; Roman & Darling, 2007).

### Trade data

There are no customs records for freshwater red algae, as they are not traded. We sought for trade data on ornamental aquatic plants and freshwater fish instead, because freshwater red algae hitchhike on those traded organisms. We found no customs records on ornamental aquatic plants imported to and exported from Taiwan in the Taiwan Customs database. But we found import and export data for ornamental freshwater fish between 2013 and 2017 (under the HS code “0301110090 Other Ornamental Fish, Freshwater” from the Customs Administration, Ministry of Finance, Taiwan; https://portal.sw.nat.gov.tw/APGA/GA03).

## RESULTS

ABGD, PTP, and GMYC inferred 27, 28, and 28 mOTUs in the 213 samples collected for this study, respectively (**Fig. S3**). We took the most conservative estimate of 27 mOTUs (26 are freshwater taxa and one is brackish) inferred using ABGD (initial partition with prior maximal distance, *p* = 7.74 x 10^−3^; distance K80 Kimura, MinSlope = 1.5). The ABGD analysis indicated that globally there are 170 mOTUs of freshwater red macroalgae, of which 26 (*15%) are found in Taiwan. A rarefaction analysis showed that excellent sampling effort was achieved to detect mOTUs (95% and 96% sample-size based completeness for the field and aquarium samples, respectively; **Fig. 3**).

**Figure 3.**
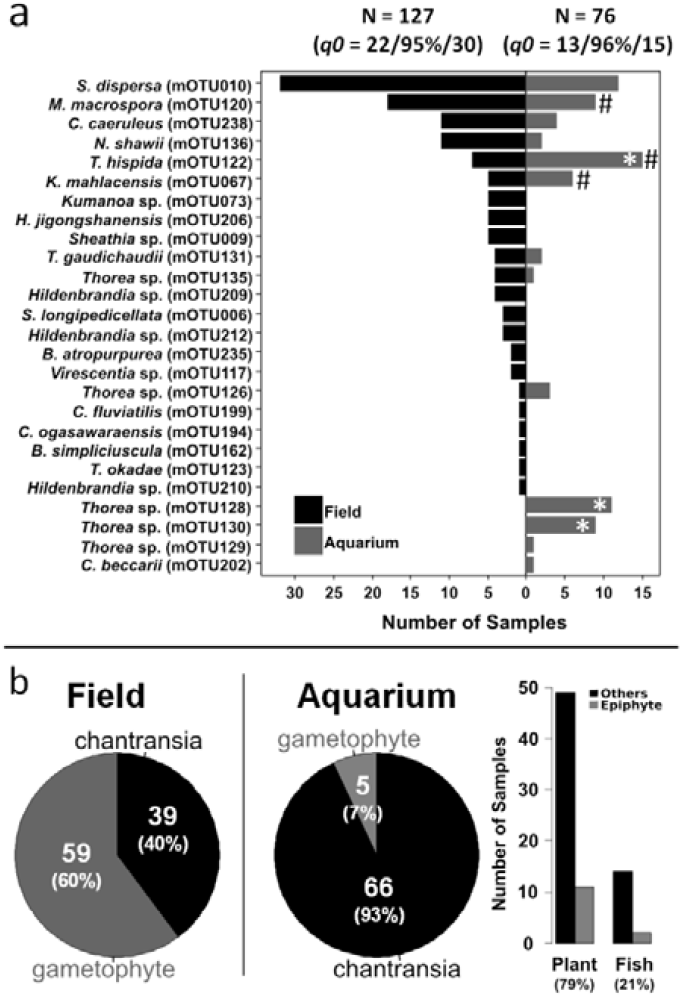
Distribution of freshwater red algal mOTUs (a) and proportion of chantransia (versus gametophyte) (b) found in the field and aquarium samples from Taiwan. Only the specimens of the taxa of Batrachospermales and Thoreales were counted for panel (b). The observed number of mOTUs, sampling effort, and the number of maximum mOTUs projected using iNEXT are shown in parentheses at the top of panel (a). The type of tank where the aquarium samples were collected is indicated in panel (b). Taxa exhibiting no genetic variation in either the field or aquarium samples are marked by the hash sign. The asterisks indicate a significant difference as per Fisher’s exact test (*p* < 0.05).

Of the 26 freshwater mOTUs, 13 (50%) are found in the field only, four (*15%) in aquaria only, and nine (*35%) mOTUs in both the field and aquaria (**Fig. 3**). We have enough samples (at least three from the field and aquaria) for only five of the mOTUs to estimate nucleotide diversity and haplotype diversity (**Table 1**). Therefore, we identified potential introduced taxa among the five mOTUs based on genetic and geographical data. Three of these five mOTUs (*Kumanoa mahlacensis* mOTU067, *Montangnia macrospora* mOTU120, and *Thorea hispida* mOTU122) exhibit no local genetic variation (i.e., in Taiwan) in the field and aquarium samples (**Table 1**; **Fig. 3a**) and are found across large geographical distances (i.e., across continents). We further examined these three mOTUs as taxa that might have been introduced into Taiwan.

**Table 1.** Maximum pairwise geographical distance, haplotype diversity (Hd), and nucleotide diversity (π) of five molecular operational taxonomic units (mOTUs) found in the aquarium shops surveyed in Taiwan in this study. These taxa have been reported in East Asia, Europe, Oceania, North America, and South America. The potential introduced taxa are bold and underlined. Hd and π were estimated for the aquarium samples and the field samples in Taiwan separately, together, and in combination with the global field samples from NCBI GenBank.

| Taxa | Distance<br>(km) | Distributional region(s) | Aquarium (Taiwan) |  |  | Field (Taiwan) |  |  | Aquarium + field (Taiwan) |  |  | Aquarium + field<br>(world, including Taiwan) |  |  |
| --- | --- | --- | --- | --- | --- | --- | --- | --- | --- | --- | --- | --- | --- | --- |
| | | | n | Hd* | $\pi$ | n | Hd* | $\pi$ | n | Hd* | $\pi$ | n | Hd* | $\pi$ |
| *C. caeruleus*<br>(mOTU238) | 25,102 | E. Asia, Oceania, Europe,<br>N. America, S. America | 3 | 0.000 (1) | 0.00000 | 11 | 0.182 (2) | 0.00171 | 14 | 0.473 (3) | 0.00219 | 72 | 0.487 (4) | 0.00167 |
| **<u>M. macrospora</u>**<br>**(mOTU120)** | 24,792 | E. Asia, S. America | 7 | 0.000 (1) | 0.00000 | 18 | 0.000 (1) | 0.00000 | 25 | 0.000 (1) | 0.00000 | 63 | 0.620 (8) | 0.01225 |
| **<u>K. mahlacensis</u>**<br>**(mOTU067)** | 16,394 | E. Asia, Europe,<br>N. America | 6 | 0.000 (1) | 0.00000 | 5 | 0.000 (1) | 0.00000 | 11 | 0.000 (1) | 0.00000 | 16 | 0.125 (2) | 0.00293 |
| **<u>T. hispida</u>**<br>**(mOTU122)** | 16,186 | E. Asia, Europe,<br>N. America | 14 | 0.000 (1) | 0.00000 | 7 | 0.000 (1) | 0.00000 | 21 | 0.000 (1) | 0.00000 | 38 | 0.149 (2) | 0.00035 |
| *S. dispersa*<br>(mOTU010) | 10,175 | E. Asia, N. America | 13 | 0.682 (4) | 0.00633 | 26 | 0.538 (3) | 0.00365 | 38 | 0.572 (5) | 0.00442 | 87 | 0.742 (7) | 0.00344 |

\*The number of distinct haplotypes is parenthesized.

For *M. macrospora* (which has decent global sampling), a haplotype network was reconstructed to identify the potential source population(s) of this introduced alga (**Fig. 4**). Eight distinct *rbc*L haplotypes were recovered (designated H1 to H8). All eight haplotypes (H1 to H8) were found in South America (specifically, Bolivia, French Guiana, and Brazil), whereas only two haplotypes (H1 and H2) were found in East Asia. H1 appears to be the most geographically widespread haplotype (at least based on the present sampling), and it is the only haplotype found in Taiwan in the field and in aquaria. Interestingly, H2 was not found in Taiwan but was found in Hong Kong, Thailand, and South America. These data suggest that South America might have been the source of *M. macrospora* (high haplotype diversity) and East Asia (with sampling densely concentrated in Taiwan in this study) might have been the sink (low haplotype diversity).

**Figure 4.**
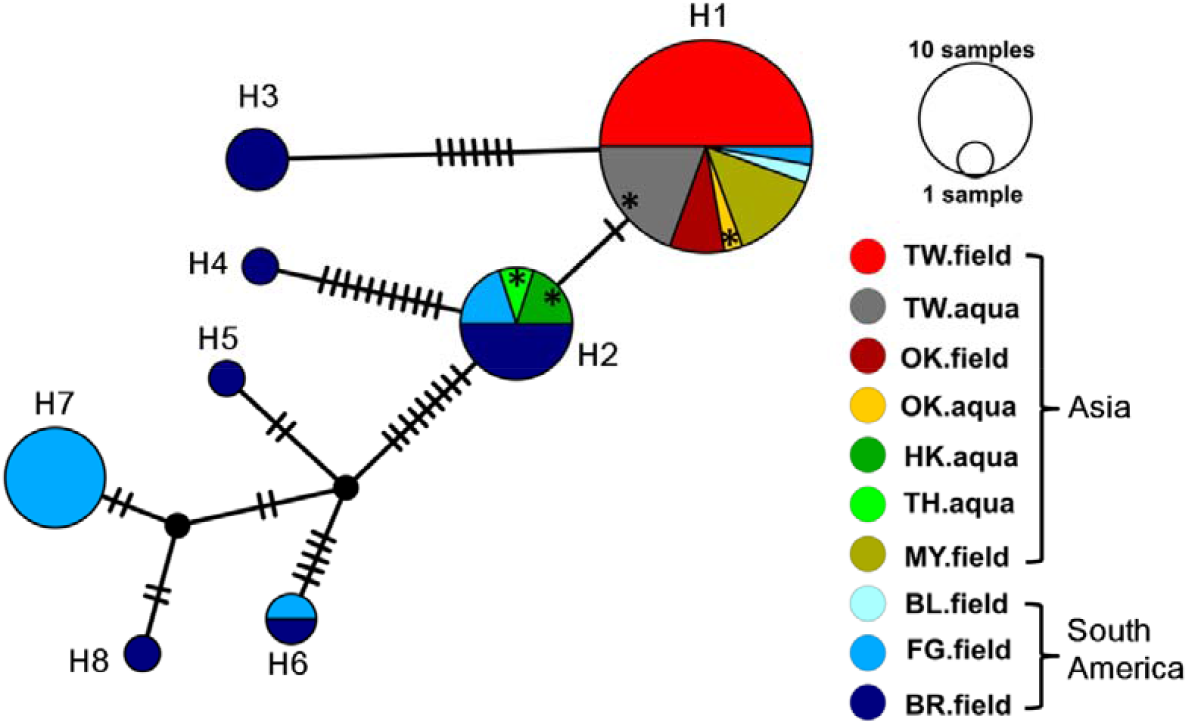
Haplotype network of *Montagnia macrospora*, a potential introduced species identified in this study. A network of eight distinct haplotypes (H1 to H8) was inferred by TCS. The hatch marks on the links between the haplotypes represent mutational steps. The asterisks indicate the aquarium samples. The field and aquarium samples from East Asia contain only H1 and H2 (low haplotype diversity), whereas all eight haplotypes were found in the field samples from South America (high haplotype diversity). Abbreviations: BL, Bolivia; BR, Brazil; FG, French Guiana; HK, Hong Kong; MY, Malaysia; OK, Okinawa; TH, Thailand; TW, Taiwan.

Of the freshwater red algae we found in aquarium shops, only the algae of Batrachospermales (*Kumanoa, Sheathia*, and *Virescentia* in our data) and Thoreales (*Nemalinopsis* and *Thorea* in our data) have morphologically distinct alternating life stages. We observed that the relative proportion of gametophyte and chantransia (which were identified morphologically) is different between the field and aquarium samples (*p* = 6.34 x 10^−12^, Fisher’s exact test; **Fig. 3b**), indicating that chantransia (the asexual stage) is the more frequently occurring reproductive form in aquaria. Chantransia do not seem to have substrate preference between aquatic plant tanks and fish tanks (*p* = 0.86, Fisher’s exact test; **Fig. 3b**). However, we observed a higher proportion of chantransia specimens in aquatic plant tanks than fish tanks (**Fig. 3b**), suggesting that chantransia might hitchhike more often on ornamental aquatic plants than animals. Also, we observed that in both types of tanks, chantransia seem to grow preferentially on inanimate substrates (e.g., plastic materials or rocks) rather than aquatic plants or animals (**Fig. 3b**; **Table S1**).

## DISCUSSION

The diversity of freshwater red algae in the aquarium trade has been largely unexplored, because the algae are easy to miss as minuscule spores and are often morphologically indistinguishable (i.e., cryptic). Collectively, only 10 species of freshwater red macroalgae have been documented in Taiwan on the basis of molecular or morphological evidence (Wu, 1999; Wu, 2001; Liu et al., 2004; Chou & Wang, 2006; Vis et al., 2010; Chou et al., 2014; Chou et al., 2015). Through our broad DNA barcode-based survey of field sites and aquarium shops across Taiwan, we found 26 mOTUs of freshwater red macroalgae (13 mOTUs in the field only, four mOTUs in aquaria only, and nine mOTUs in the field and aquaria). To the extent that the mOTUs correspond to distinct species, this result indicates that the biodiversity of freshwater red macroalgae in Taiwan is substantially richer than previously reported.

In this study, we found that some freshwater red algae might have been introduced via the aquarium trade, in line with previous observations (e.g., Stoyneva et al., 2006; Kato et al., 2009). Using the amount of genetic variation and the extent of geographical distribution as criteria, we identified three mOTUs (*Kumanoa mahlacensis, Montangnia macrospora*, and *Thorea hispida)* as potential introduced species among the 13 mOTUs found in aquaria in Taiwan. The three mOTUs were also found in the field samples, suggesting successful introduction. *M. macrospora* has already been reported to be an alien species in a eutrophic artificial dam in Okinawa, Japan (Kato et al., 2009). *T. hispida* has been reported to be cosmopolitan with low genetic diversity, possibly partly due to the aquarium trade (Johnston et al., 2018). This epizoic alga can hitchhike on rusty crayfish (*Orconectes rusticus)*, which is a common invasive species in Asia (Fuelling et al., 2012). As far as we know, *K. mahlacensis* is not known as an introduced species elsewhere in the world. On the basis on our data, we hypothesize that *K. mahlacensis* has been introduced to Taiwan via the global aquarium trade.

Routes of introduction can be inferred based on genetic and occurrence data (e.g., Bonett et al., 2007; Muirhead et al., 2008; Le Roux et al., 2011). We illustrate this using *M. macrospora* as an example. Kato et al. (2009) has established that *M. macrospora* was introduced by the aquarium trade between South America and Okinawa, Japan. Later, this alga was reportedly found in the field in Taiwan and Malaysia (Chou et al., 2014; Johnston et al., 2014). In our study, *M. macrospora* was found in the aquarium shops surveyed in Taiwan, Okinawa (Japan), Hong Kong, and Thailand. We observed that the nucleotide diversity and haplotype diversity of the *rbc*L sequences from the South America samples of *M. macrospora* (n = 25, π = 0.02183, Hd = 0.820, 8 haplotypes; from Bolivia, Brazil, and French Guiana) are higher than the nucleotide diversity and haplotype diversity of the sequences from the East Asia samples (n = 38, π = 0.00004, Hd = 0.149, 2 haplotypes; from Taiwan, Thailand, Okinawa in Japan, and Malaysia). Only two haplotypes (H1 and H2) were found in East Asia (**Fig. 4**). Interestingly, the trade data of ornamental fish in Taiwan indicate that there is high import activity to Taiwan from South America (**Fig. S4**; **Table S2**) and therefore that an introduction route between the two regions is plausible. These data suggest that *M. macrospora* might have been introduced to East Asia (possibly more than once, given that there are two haplotypes in East Asia) from South America (but the country of origin is unknown due to limited field sampling) and then it may spread further among the regions in East Asia through the aquarium trade. Plausible introduction routes for the other two cases (*K. mahlacensis* and *T. hispida*), however, are less clear because of insufficient sampling from field populations (**Table 1**).

In addition to potential introduced taxa, we identified cryptic ones. Our analysis uncovered three cryptic taxa of *Thorea* (mOTU128, mOTU129, and mOTU130) (**Fig. 3**; **Fig. S3**). These cryptic taxa may be novel species that have not yet been circumscribed taxonomically. Interestingly, two of them (mOTU128 and mOTU130) were found to be common in the aquarium shops in Taiwan but not in the field anywhere in the world (at least based on the current sampling of *rbc*L sequences) (**Fig. 3**; **Fig. S3**). This observation raises the intriguing possibility that the aquarium trade can act as a reservoir for some algal species which may not be widespread in the field or may be even endangered (see below). These mOTUs are examples of cryptic taxa that have gone unnoticed under the current morphology-based biodiversity monitoring programs. For example, chantransia (the sporophytic stage) of freshwater red macroalgae are minute hitchhikers and morphologically alike. Such cryptic taxa have the potential to become invasive. Therefore, it is important to incorporate better tools, such as DNA barcodes, into biodiversity monitoring approaches.

Besides introduced taxa and cryptic taxa, the aquarium trade may preserve endangered freshwater red algae. For example, *Nemalionopsis shawii* (previously named *N. tortuosa*) and *Thorea gaudichaudii* are regarded as endangered in Japan, possibly due to anthropogenic disturbances (Brodie et al., 2009). Consistent with this, we preliminarily did not find these algae in the four aquarium samples that we collected in Okinawa, Japan (**Table S1**). However, we did find a few uncommon instances of *N. shawii* (mOTU136; in 10 field samples and two aquarium samples) and *T. gaudichaudii* (mOTU131; in four field samples and two aquarium samples) in Taiwan (**Fig. 3**), likely because of a much greater sampling effort (in total, 127 field samples and 76 aquarium samples were collected in Taiwan). If endangered freshwater red algae were accidentally transferred between the field and aquaria (which may be the case in Taiwan), then a wide survey of aquaria in Japan may reveal *N. shawii* and *T. gaudichaudii.* Thus, the aquarium trade may act as an *ex situ* preservation site for *N. shawii, T. gaudichaudii*, and probably other freshwater algae, which are being threatened by water pollution caused by urbanization or industrialization (Sheath & Hambrook, 1990; Sheath & Vis, 2015).

The criteria adopted in this study to identify potential introduced taxa may be too strict. Introduction of a species into a non-native habitat can cause a genetic bottleneck that results in a drastic reduction of genetic diversity; however, multiple introductions of a species into the same non-native habitat can lead to appreciable levels of genetic diversity (reviewed in Wilson et al., 2009). Thus, potential cases of introduction elsewhere (or in Taiwan) may be missed under our stringent criteria. One example is *Composopogon caeruleus*, which is frequently found in aquaria (e.g., Stoyneva et al., 2006; Levent & Bud, 2010; Carlile & Sherwood, 2013; Necchi Jr. et al., 2013). This alga has been reported as a tropical invader in Belgium via the aquarium trade (Stoyneva et al., 2006). However, we ruled out this alga as an introduced species in Taiwan because it did not meet our criterion of no local nucleotide or haplotype diversity (**Table 1**).

For this study, we selected *rbc*L as a marker to track potential introductions because the sequence data for this gene are abundantly available and well represented geographically for freshwater red algae compared to other nuclear, mitochondrial, and plastid loci. The *rbc*L sequences of the specimens sampled worldwide enabled us to identify tentative cases of freshwater red algae introduced to Taiwan from distant regions and vice versa. In future investigations, it may be fruitful to explore the utility of alternative marker genes, such as *rpoC1* (Zhan et al., 2020), and population-level markers, such as microsatellite loci (e.g., Kinziger et al., 2011), to unveil candidate algal introductions alongside *rbc*L. Population-level markers would help to better evaluate differences in genetic diversity between an introduced population and its potential source population(s) (e.g., Muirhead et al., 2008), thereby refining our criteria of potential introduced algae. Presently, a disadvantage of using these marker loci is their geographically restricted sampling, which limits our capacity to detect long-distance introductions (e.g., between Taiwan and South America). Enough sequence data for those marker loci (with geographically broad sampling) need to be obtained before employing the marker loci to detect potential instances of introduced freshwater red macroalgae.

Here, we analyzed *rbc*L sequences for evidence that freshwater red macroalgae are dispersed via the global aquarium trade. Besides the aquarium trade, waterfowl may serve as another mechanism of short- or long-distance dispersal of freshwater red algae. Waterfowl are thought to be active dispersers of aquatic alien organisms, including algae (reviewed in Reynolds et al., 2015). Short-distance dispersal of freshwater red algae (such as organisms of Batrachospermales) by waterfowl is plausible. We found one case of potential introduction via short-distance dispersal – *N. shawii*, which is found only in East Asia based on the present sampling. However, long-distance, transoceanic dispersal of the freshwater algae by waterfowl is presumably unlikely, because these algae obviously cannot tolerate salt water or long-term desiccation. In this study, examples of freshwater algae dispersed over the oceans are *M. macrospora* and *T. hispidia*. Our genetic diversity estimates suggest that *M. macrospora* has been introduced from South America to East Asia, and indicated that *T. hispidia* is distributed in East Asia, Europe, and North America. Hence, for these two algae, the aquarium trade is a more probable mechanism of dispersal than waterfowl. Moreover, another possible mechanism to introduce alien freshwater algae is ballast water discharge, which is known to wreck environmental havoc by spreading invasive species (e.g., zebra mussel). Manny et al. (1991) speculated that *C. caeruleus*, a freshwater red alga, might be transported via ballast water discharge, but so far it remains to be a speculation. Although ballast water discharge is an unlikely mechanism to introduce freshwater algae into Taiwan (an island surrounded by seawater), it may transport freshwater algae between distant locations connected by freshwater bodies, such as lakes and rivers, in other parts of the world.

DNA barcoding of individual specimens, as performed in this study, can be labor-intensive and time-consuming, and it provides a restricted view of the biodiversity in an ecosystem. A powerful alternative is environmental DNA metabarcoding enabled by high-throughput sequencing. Recently, this approach has been applied for large-scale biodiversity monitoring of various ecosystems (reviewed in Thomsen & Willerslev, 2015). We envisage that environmental DNA metabarcoding can accelerate the identification of introduced, cryptic, and endangered taxa of freshwater red algae in the field and aquaria, thereby expanding our knowledge of the biodiversity of these algae in the wild and artificial environments.

Hitchhiking freshwater red algae have gone overlooked over the past decades. We surveyed the biodiversity of freshwater red macroalgae in the field and aquaria across Taiwan and in nearby regions. We identified potential introductions of freshwater red macroalgae into Taiwan through the global aquarium trade. We anticipate that our data will serve as a taxonomic resource for future large-scale monitoring programs that utilize DNA barcoding and environmental DNA metabarcoding, especially around large urban centers where aquarium trade activity is higher (Strechker et al., 2011). Overlooked hitchhikers, such as freshwater red macroalgae, are not regulated by CITES (Convention on International Trade in Endangered Species of Wild Fauna and Flora), hampering the prevention of their spread via the aquarium trade. It is important to educate aquarists and the public about the proper disposal of aquarium waste (e.g., putting it in solid waste for compost, microwaving/freezing it prior to the waste disposal, or treating it with chemicals), as suggested by Padilla & Williams (2004), Patoka et al. (2016), and Vranken et al. (2018). Furthermore, detailed studies about the potential ecological and social concerns of algal introductions (e.g., the homogenization of global taxonomic diversity, loss of local biodiversity, and potential transfer of pests or pathogens) are needed to better determine whether or not hitchhikers should be actively controlled or eradicated in the aquarium trade.

## ACKNOWLEDGEMENTS

We thank Dr. Jaruwan Mayakun (Prince of Songkla University) and Dr. Paul Geraldino (University of San Carlos) for helping to collect the specimens from Thailand and the Philippines. We also thank Prof. Sarah P. Otto (University of British Columbia) for critically reviewing the manuscript. This work was supported by grants from Ministry of Science and Technology, Taiwan to SLL (MOST108-2621-B-029-005-MY3).

## DATA AVAILABILITY STATEMENT

The *rbc*L sequences obtained in this study are deposited in GenBank under the accession numbers: MH835465-MH835677.

## Author contributions

S.H.Z. and S.L.L. conceived the idea. S.L.L. and T.Y.H. designed the study. S.L.L., S.H.Z., T.Y.H., and S.S. performed specimen collection and laboratory experiment. S.L.L., L.Y., and T.C.K. analyzed the data. S.H.Z. and S.L.L. wrote the manuscript. All authors discussed the results and reviewed the manuscript.

## REFERENCES

Ahrens, D., Fujisawa, T., Krammer, H.-J., Eberle, J., Fabrizi, S., & Vogler, A. P. (2016). Rarity and incomplete sampling in DNA-based species delimitation. Systematic Biology, 65(3), 478–494.

Allendorf, F. W., & Lundquist, L. L. (2003). Introduction: population biology, evolution, and control of invasive species. Conservation Biology, 17(1), 24–30.

Armstrong, K. F., & Ball, S. L. (2005). DNA barcodes for biosecurity: invasive species identification. Philosophical Transactions of the Royal Society B: Biological Sciences, 360(1462), 1813–1823.

Barrett, S. C. H., & Husband, B. C. (1990). The genetics of plant migration and colonization. In Plant Population Genetics, Breeding, and Genetic Resources (eds. Brown, A. H. D.; Clegg, M. T.; Kahler, A. L.; Weir, B. S.), pp. 254–277. Sinauer Associates, Sunderland, Massachusetts, USA.

Bonett, R. M., Kozak, K. H., Vieites, D. R., Bare, A., Wooten, J. A., & Trauth, S. E. (2007). The importance of comparative phylogeography in diagnosing introduced species: a lesson from the seal salamander, *Desmognathus monticola*. BMC Ecology, 7, 7.

Carlile, A. L. & Sherwood, A. R. (2013). Phylogenetic affinities and distribution of the Hawaiian freshwater red algae (Rhodophyta). Phycologia, 52, 309–319.

Carstens, B. C., Pelletier, T. A., Reid, N. M., & Satler, J. D. (2013). How to fail at species delimitation. Molecular Ecology, 22(17), 4369–4383.

Chou, J.-Y, & Wang, W.-L. (2006). *Batrachospermum arcuatum* Kylin (Batrachospermales, Rhodophyta), a freshwater red alga newly recorded in Taiwan. Taiwania, 51(1), 58–63.

Chou, J.-Y, Wen, Y-D., & Wang, W.-L. (2014). Morphological and molecular data confirm new records of three freshwater red algae, *Batrachospermum macrosporum, Nemalionopsis tortuosa* and *Caloglossa leprieurii* in Taiwan. Nova Hedwigia, 98(1), 233–246.

Chou, J.-Y., Liu, S.-L., Wen, Y-D., & Wang, W.-L. (2015). Phylogenetic analysis of *Bangiadulcis atropurpurea* (A. Roth) W. A. Nelson and *Bangia fuscopurpurea* (Dillwyn) Lyngbye (Bangiales, Rhodophyta) in Taiwan. Archives of Biological Science, Belgrade, 67(2), 445–454.

Diaz Tapia, P., Maggs, C. A., Macaya, E. C., & Verbruggen, H. (2018). Widely distributed red algae often represent hidden introductions, complexes of cryptic species or species with strong phylogeographic structure. Journal of Phycology 54(6), 829–839.

Dlugosch, K. M., & Parker, I. M. (2008). Founding events in species invasions: genetic variation, adaptive evolution, and the role of multiple introductions. Molecular Ecology, 17, 431–449.

Duggan, I. C., & Pullan, S. G. (2017). Do freshwater aquaculture facilities provide an invasion risk for zooplankton hitchhikers? Biological Invasions, 19(1), 307–314.

Duggan, I. C., Champion, P. D., & MacIsaac, H. J. (2018). Invertebrates associated with aquatic plants bought from aquarium stores in Canada and New Zealand. Biological Invasions, 20(11): 3167–3178.

Dupuis, J. R., Roe, A. D., & Sperling, F. A. (2012). Multi-locus species delimitation in closely related animals and fungi: one marker is not enough. Molecular Ecology, 21(18), 4422–4436.

Edgar, R. C. (2004). MUSCLE: multiple sequence alignment with high accuracy and high throughput. Nucleic Acids Research, 32(5),1792–1797.

Edgar, R. C. (2010). Search and clustering orders of magnitude faster than BLAST. Bioinformatics, 26(19), 2460–2461.

Esselstyn, J. A., Evans, B. J., Sedlock, J. L., Anwarali Khan, F. A., & Heaney, L. R. (2012). Single-locus species delimitation: a test of the mixed Yule-coalescent model, with an empirical application to Philippine round-leaf bats. Proceedings of the Royal Society B: Biological Sciences, 279(1743), 3678–3686.

Fuelling, L. J., Adams, J. A.; Badik, K. J., Bixby, R. J., Caprette, C. L., Caprette, H. E., Chiasson, W. B., Davies, C. L., Decolibus, D. T., Glascock, K. I., Hall, M. M., Perry, W. L., Schultz, E. R., Taylor, D. A., Vis, M. L., & Verb, R. G. (2012). An unusual occurrence of *Thorea hispida* (Thore) Desvaux chantransia on rusty crayfish in West Central Ohio. Nova Hedwigia, 94(3-4), 355–366.

Guiry, M. D., & Guiry, G. M. (2020). AlgaeBase. World-wide electronic publication, National University of Ireland, Galway. https://www.algaebase.org; searched on 07 March 2020.

Hall, T. A. (1999). BioEdit: A user-friendly biological sequence alignment editor and analysis program for Windows 95/98/NT. Nucleic Acids Symposium Series, 41, 95–98.

Hawes, I., Howard-Williams, C., Wells, R. D. S., & Clayton, J. S. (1991). Invasion of water net, *Hydrodictyon reticulatum:* the surprising success of an aquatic plant new to our flora. New Zealand Journal of Marine and Freshwater Research, 25(3), 227–229.

Hsieh, T. C., Ma, K. H., & Chao, A. (2014). iNEXT: An R package for interpolation and extrapolation in measuring species diversity. (http://chao.stat.nthu.edu.tw/blog/software-download/inext-r-package/).

Johnston, E. T., Dixon, K. R., West, J. A., Buhari, N., & Vis, M. L. (2018). Three gene phylogeny of the Thoreales (Rhodophyta) reveals high species diversity. Journal of Phycology, 54(2), 159–170.

Johnston, E. T., Lim, P.-E., Buhari, N., Keil, E. J., Djawad, M. I., & Vis, M. L. (2014). Diversity of freshwater red algae (Rhodophyta) in Malaysia and Indonesia from morphological and molecular data. Phycologia, 53(4), 329–341.

Kaštovský, J., Hauer, T., Mareš, J., Krautová, M., Beša, T., Komárek, J., Desirtivá, B., Heteša, J., Hindáková, A., Houk, V., Janeček, E., Kopp, R., Marvan, P., Pumann, P., Skácelová, O., & Zapomělová, E. (2010). A review of the alien and expansive species of freshwater cyanobacteria and algae in the Czech Republic. Biological Invasions, 12(10), 3599–3625.

Kato, A., Morita, N., Hiratsuka, T., & Suda, S. (2009). Recent introduction of a freshwater red alga Chantransia macrospora (Batrachospermales, Rhodophyta) to Okinawa, Japan. Aquatic Invasions, 4(4), 567–574.

Kaufmann, B. (2010). Algen-Fibel Aquarium: Kein Problem mit Süßwasseralgen (pp. 94). Germany: Dähne Verlag Press.

Kinziger, A. P., Nakamoto, R. J., Anderson, E. C., & Harvey, B. C. (2011). Small founding number and low genetic diversity in an introduced species exhibiting limited invasion success (speckled dace, *Rhinichthys osculus)*. Ecology and Evolution, 1(1), 73–84.

Knowles, L. L., & Carstens, B. C. (2007). Delimiting species without monophyletic gene trees. Systematic Biology, 56(6), 887–895.

Le Roux, J. J., Brown, G. K., Byrne, M., Ndlovu, J., Richardson, D. M., Thompson, G. D., & Wilson, J. R. U. (2011). Phylogeographic consequences of different introduction histories of invasive Australian *Acacia* species and Paraserianthes lophantha (Fabaceae) in South Africa. Diversity and Distributions, 17(5), 861–871.

Leigh, J. W., & Bryant, D. (2015). PopART: Full-feature software for haplotype network construction. Methods in Ecology and Evolution, 6(9), 1110–1116.

Leliaert, F., Verbruggen, H., Vanormelingen, P., Steen, F., López-Bautista, J. M., Zuccarello, G. C., & De Clerck, O. (2014). DNA-based species delimitation in algae. European Journal of Phycology, 49(2), 179–196.

Lepage, T., Bryant, D., Philippe, H., & Lartillot, N. (2007). A general comparison of relaxed molecular clock models. Molecular Biology and Evolution, 24(12), 2669–2680.

Levente, V., & Bud, I. (2010). Effects of hydrogen peroxide on Compsopogon caeruleus (Rhodophycophyta) and two superior plants. AACL Bioflux, 3(5), 367–372.

Lin, C. K., & Blum, J. L. (1977). Recent invasion of a red alga (*Bangia atropurpurea)* in Lake Michigan. Journal of the Fisheries Research Board of Canada, 34(12), 2413–2416.

Lin, S.-M., Fredericq, S., & Hommersand, M. H. (2001). Systematics of the Delesseriaceae (Ceramiales, Rhodophyta) based on LSU rDNA and *rbc*L sequences, including the Phycodryoideae, subfam. nov. Journal of Phycology, 37(5), 881–899.

Liu, S.-L., Wang, L.-C., & Wang, W.-L. (2004). Inorganic carbon utilization of the freshwater red alga *Compsopogon coeruleus* (Balbis) Montagne (Compsopogonaceae, Rhodophyta) evaluated by *in situ* measurement of chlorophyll fluorescence. Taiwania, 49(3), 207–217.

Luther, H. (1979). *Chara conniens* in the Baltic sea area. Annales Botanici Fennici, 16, 141–150.

Maddison, W. P., & Maddison, D. R. (2011). Mesquite: a modular system for evolutionary analysis. Version 2.75. http://mesquiteproject.org.

Manny, B. A., Esdall, T., & Wujek, D. (1991). *Compsopogon* cf. *coeruleus*, a benthic red alga (Rhodophyta) new to the Laurentian Great Lakes. Canadian Journal of Botany, 69(6), 1237–1240.

Muirhead, J. R., Gray, D. K., Kelly, D. W., Ellis, S. M., Heath, D. D., & Macisaac, H. J. (2008). Identifying the source of species invasions: sampling intensity vs. genetic diversity. Molecular Ecology, 17, 1020–1035.

Novak, S. J., & Mack, R. N. (1993). Genetic variation in *Bromus tectorum* (Poaceae): comparison between native and introduced populations. Heredity, 71, 167–176.

Novak, S. J., & Mack, R. N. (2005). Genetic bottlenecks in alien plant species: influences of mating systems and introduction dynamics. In: Species Invasions: Insights into Ecology, Evolution, and Biogeography (eds. Sax, D. F., Stachowicz, J. J., Gaines, S. D.), pp. 201–228. Sinauer Associates, Sunderland, Massachusetts, USA.

Necchi Jr., O., Garcia Fo, A. S., Salomaki, E. D., West, J. A., Aboal, M., & Vis, M. L. (2013). Global sampling reveals low genetic diversity within *Compsopogon* (Composopogonales, Rhodophyta). European Journal of Phycology, 48(2), 152–162.

Padilla, D. K., & Williams, S. L. (2004). Beyond ballast water: aquarium and ornamental trades as sources of invasive species in aquatic ecosystems. Frontiers in Ecology and the Environment, 2(3), 131–138.

Patoka, J., Bláha, M., Kalous, L., Vrabec, V., Buřič, M., & Kouba, A. (2016). Potential pest transfer mediated by international ornamental plant trade. Scientific Reports, 6, 25896.

Pecnikar, Z. F., & Buzan, E. V. (2014). 20 years since the introduction of DNA barcoding: from theory to application. Journal of Applied Genetics, 55(1), 43–52.

sPons, J., Barraclough, T. G., Gomez-Zurita, J., Cardoso, A., Duran, D. P., Hazell, S., Kamoun, S., Sumlin, W. D., & Vogler, A. P. (2006). Sequence-based species delimitation for the DNA taxonomy of undescribed insects. Systematic Biology, 55(4), 595–609.

Puillandre, N., Lambert, A., Brouillet, S., & Achaz, G. (2012). ABGD, Automatic Barcode Gap Discovery for primary species delimitation. Molecular Ecology, 21(8), 1864–1877.

Rahel, F. J. (2007). Biogeographic barriers, connectivity and homogenization of freshwater faunas: it’s a small world after all. Freshwater Biology, 52(4), 696–710.

Reynolds, C., Miranda, N. A., & Cumming, G. S. (2015). The role of waterbirds in the dispersal of aquatic alien and invasive species. Diversity and Distributions, 21(7), 744–754.

Riedel, A., Sagata, K., Suhardjono, Y. R., Tänzler, R., & Balke, M. (2013). Integrative taxonomy on the fast track – towards more sustainability in biodiversity research. Frontiers in Zoology, 10, 15.

Roman, J., & Darling, J. A. (2007). Paradox lost: genetic diversity and the success of aquatic invasions. Trends in Ecology and Evolution, 22(9), 454–464.

Ronquist, F., Teslenko, M., van der Mark, P., Ayres, D. L., Darling, A., Höhna, S., Larget, B., Liu, L., Suchard, M. A., & Huelsenbeck, J. P. (2012). MrBayes 3.2: efficient Bayesian phylogenetic inference and model choice across a large model space. Systematic Biology, 61(3), 539–542.

Rozas, J., Ferrer-Mata, A., Sánchez-DelBarrio, J. C., Guirao-Rico, S., Librado, P., Ramos-Onsins, S. E., & Sánchez-Gracia, A. (2017). DnaSP v6: DNA sequence polymorphism analysis of large datasets. Molecular Biology and Evolution, 34(12), 3299–3302.

Schloesser, D. W., Hudson, P. L., & Nichols, S. J. (1986). Distribution and habitat of *Nitellopsis obtusa* (Characeae) in the Laurentian Great Lakes. Hydrobiologia, 133(1), 91–96.

Sheath, R. G., & Hambrook, J. A. (1990). Freshwater ecology. In K. M. Cole & R. G. Sheath (eds.), Biology of the Red Algae (pp. 423–453). New York, USA: Cambridge University Press.

Sheath, R. G., & Vis, M. L. (2015). Red Algae. In J. D. Wehr, R. G. Sheath, & J. P. Kociolek (Eds), Freshwater Algae of North America (pp. 237–264). San Diego, CA, USA: Elsevier Inc. Press.

Simberloff, D. (2014). Biological invasions: What’s worth fighting and what can be won? Ecological Engineering, 65(2014), 112–121.

Simberloff, D., Martin, J. L., Genovesi, P., Maris, V., Wardle, D. A., Aronson, J., Courchamp, F., Galil, B., García-Berthou, E., Pascal, M., Pyšek, P., Sousa, R., Tabacchi, E., & Vilà, M. (2013). Impacts of biological invasions: what’s what and the way forward. Trends in Ecology and Evolution, 28(1), 58–66.

Stoyneva, M., Vanhoutte, K., & Vyverman, W. (2006). First record of the tropical invasive alga *Compsopogon coeruleus* (Balbis) Montagne (Rhodophyta) in Flanders (Belgium). In: N. Ognjanova-Rumenova & K. Manoylov (eds.), Advances in Phycological Studies: Festschrift in Honour of Prof. Dobrina Temniskova-Topalova (pp. 203–212). PENSOFT Publishers & University Publishing House Sofia–Moscow.

Strayer, D. L. (2010). Alien species in fresh waters: ecological effects, interactions with other stressors, and prospects for the future. Freshwater Biology, 55(S1), 152–174.

Strechker, A. L., Campbell, P. M., & Olden, J. D. (2011). The aquarium trade as an invasion pathway in the Pacific Northwest. Fisheries, 36(2), 74–85.

Tamura, K., Stecher, G., Peterson, D., Filipski. A., & Kumar, S. (2013). MEGA6: Molecular evolutionary genetic analysis version 6.0. Molecular Biology and Evolution, 30(12), 2725–2729.

Thomsen, P. F., & Willerslev, E. (2015). Environmental DNA – An emerging tool in conservation for monitoring past and present biodiversity. Biological Conservation, 783, 4–18.

Vis, M. L., Feng, J., Chiasson, W. B., Xie, S. L., Stancheva, R., Entwisle, T. J., Chou, J. Y., & Wang, W. L. (2010). Investigation of the molecular and morphological variability in *Batrachospermum arcuatum* (Batrachospermales, Rhodophyta) from geographically distant locations. Phycologia, 49(6), 545–553.

Vis, M. L., Saunders, G. W., Sheath, R. G., Dunse, K., & Entwisle, T. J. (1998). Phylogeny of the Batrachospermales (Rhodophyta) inferred from rbcL and 18S ribosomal DNA gene sequences. Journal of Phycology, 34(2), 341–350.

Vranken, S., Bosch, S., Peña, V., Leliaert, F., Mineur, F., & De Clerck, O. (2018). A risk assessment of aquarium trade introductions of seaweed in European waters. Biological Invasions, 20(5), 1171–1187.

Wang, W. L., Liu, S. L., & Lin, S. M. (2005). Systematics of the calcified genera of the Galaxauraceae (Nemaliales, Rhodophyta) with an emphasis on Taiwan species. Journal of Phycology, 41(3), 685–703.

Wells, R. D. S., & Clayton, J. S. (2001). Ecological impacts of water net (*Hydrodictyon reticulatum)* in Lake Aniwhenua. New Zealand. New Zealand Journal of Ecology, 25(2), 55–63.

Wells, R. D. S., Hall, J. A., Clayton, J. S., Champion, P. D., Payne, G. W., & Hofstra, D. E. (1999). The rise and fall of water net (*Hydrodictyon reticulatum)* in New Zealand. Journal of Aquatic Plant Management, 37(2), 49–55.

Wilson, J. R. U., Dormontt, E. E., Prentis, P. J., Lowe, A. J., & Richardson, D. M. (2009). Something in the way you move: dispersal pathways affect invasion success. Trends in Ecology and Evolution, 24(3), 136–144.

Wu, J. T. (1999). Occurrence of four freshwater rhodophytes in Taiwan. Taiwania, 44(1), 145–153.

Wu, J. T. (2001). Supplements to the freshwater rhodophytes in Taiwan. Taiwania, 46(4), 359–362.

Zhan, S. H., Shih, C. C., & Liu, S. L. (2020). Reappraising plastid markers of the red algae for phylogenetic community ecology in the genomic era. Ecology and Evolution, 10(3), 1299–1310.

Zhang, J., Kapli, P., Pavlidis, P., & Stamatakis, A. (2013). A general species delimitation method with applications to phylogenetic placements. Bioinformatics, 29(22), 2869–2876.

