## Supplemental figures 1-4 for "Hidden introductions of freshwater red algae via the aquarium trade exposed by DNA barcodes"

**Supplementary Materials**

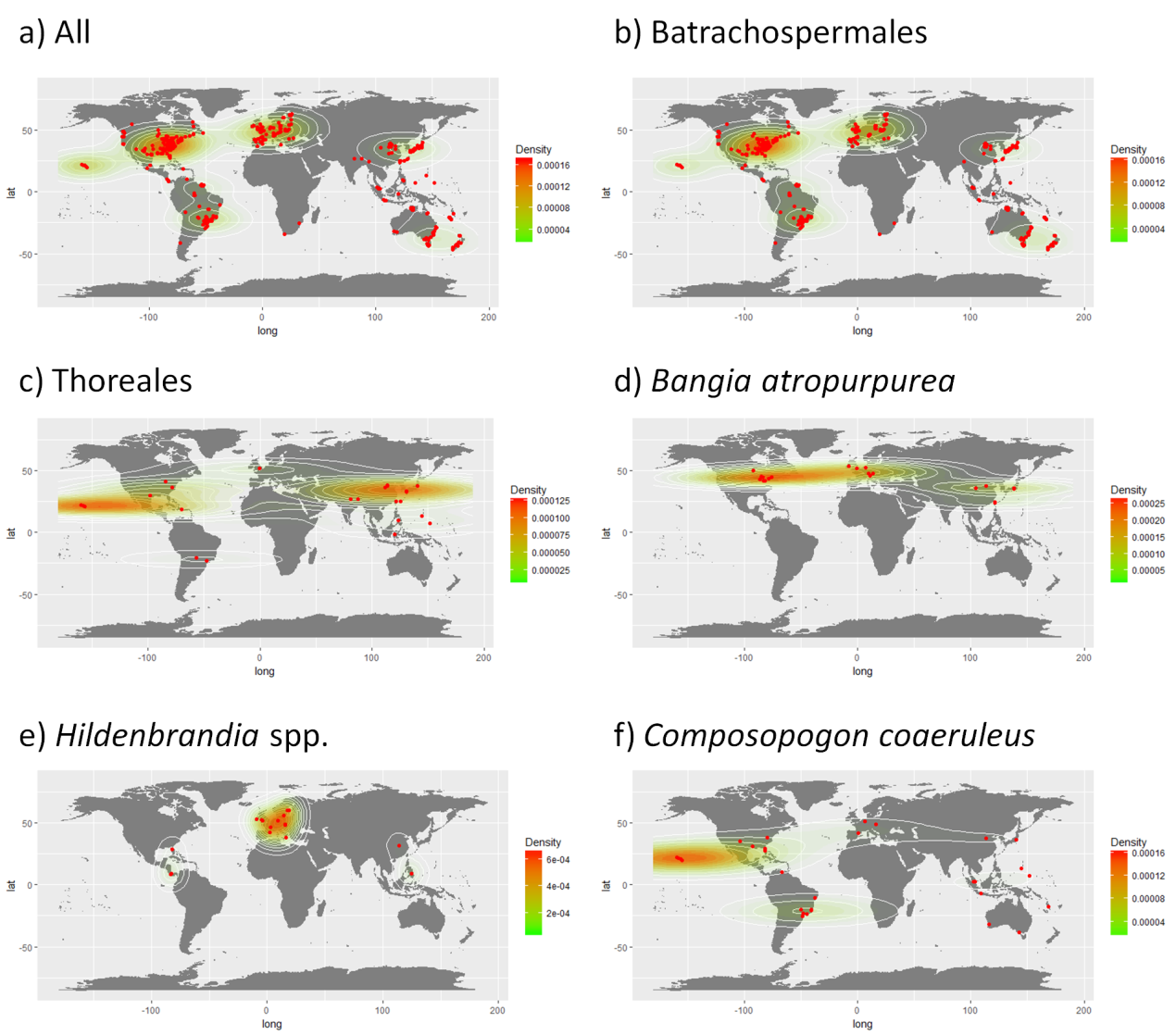

**Figure S1**. Global sampling of the GenBank records of the freshwater red macroalgae examined in this study: all taxa (a), Batrachospermales (e.g., *Batrachospermum*, *Kumanoa*, *Lemanea*, *Sheathia*, and *Virescentia*; see Table S1) (b), Thoreales (*Nemalinopsis* and *Thorea*) (c), *Bangia atropurpurea* (d), *Hildenbrandia* spp. (e), and *Compsopogon caeruleus* (f). The contours indicate the latitudinal and longitudinal density of the GenBank records (represented as red dots). It should be noted the contours do not indicate the geographical distribution of the freshwater red algae taxa.

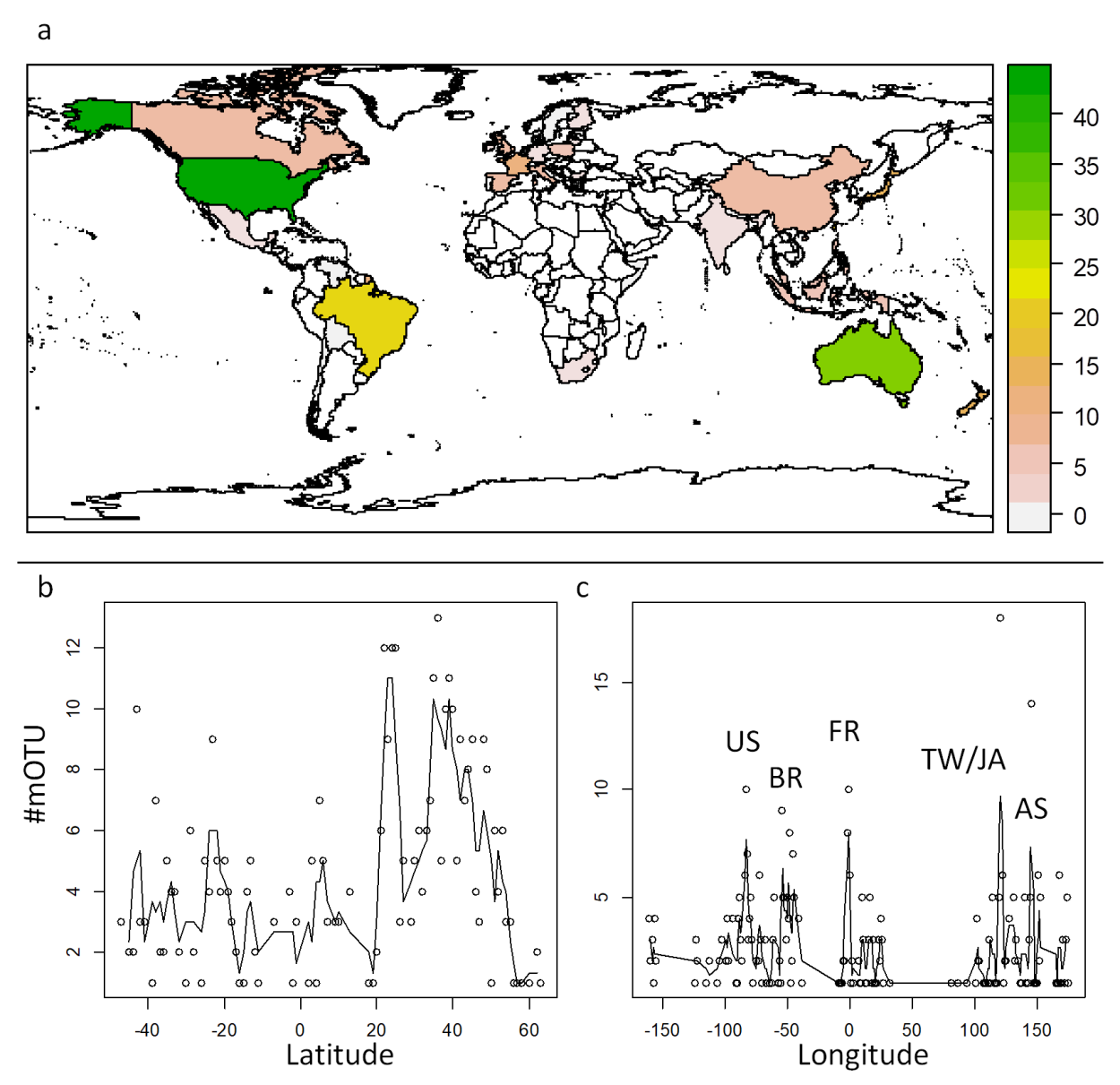

**Figure S2**. Geographical distribution of freshwater red macroalgae in different countries (a), along the latitude (b), and along the longitude (c) based on GenBank records. The number of molecular operational taxonomic units (mOTUs) was inferred using automated barcode gap discovery (ABGD).

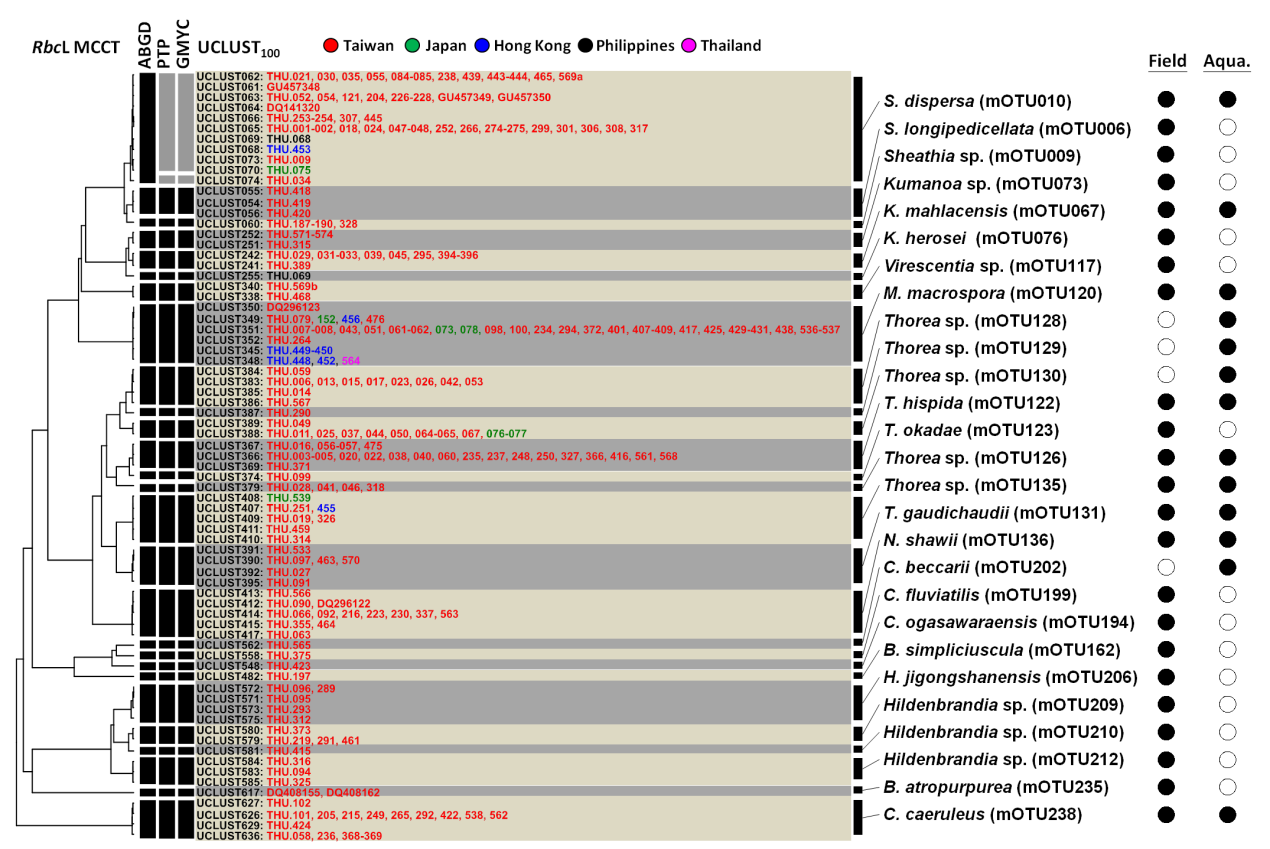

**Figure S3.** Phylogeny and diversity of the freshwater red macroalgal mOTUs in the field and aquarium samples in this study. The mOTUs were inferred using three different species delimitation methods: ABGD, PTP, and GMYC. The full set of *rbc*L sequences were clustered using USEARCH (designated as “UCLUST<number>”) before the determination of mOTUs (designated as “*Genus species* mOTU<number>”). The identification codes of the samples are colored by the geographical region where the samples were collected. The legend on the right indicates whether the samples in the UCLUST clusters were collected from the field and/or aquarium shops (black dot denotes “yes”). Abbreviations: ABGD, automated barcode gap discovery; Aqua., aquarium; GMYC, general mixed Yule-Coalescent model; MCCT, maximum clade credibility tree; mOTU, molecular operational taxonomic unit; PTP, Poisson tree process; UCLUST_100_, OTU clustering based on 100% sequence similarity threshold using USEARCH.

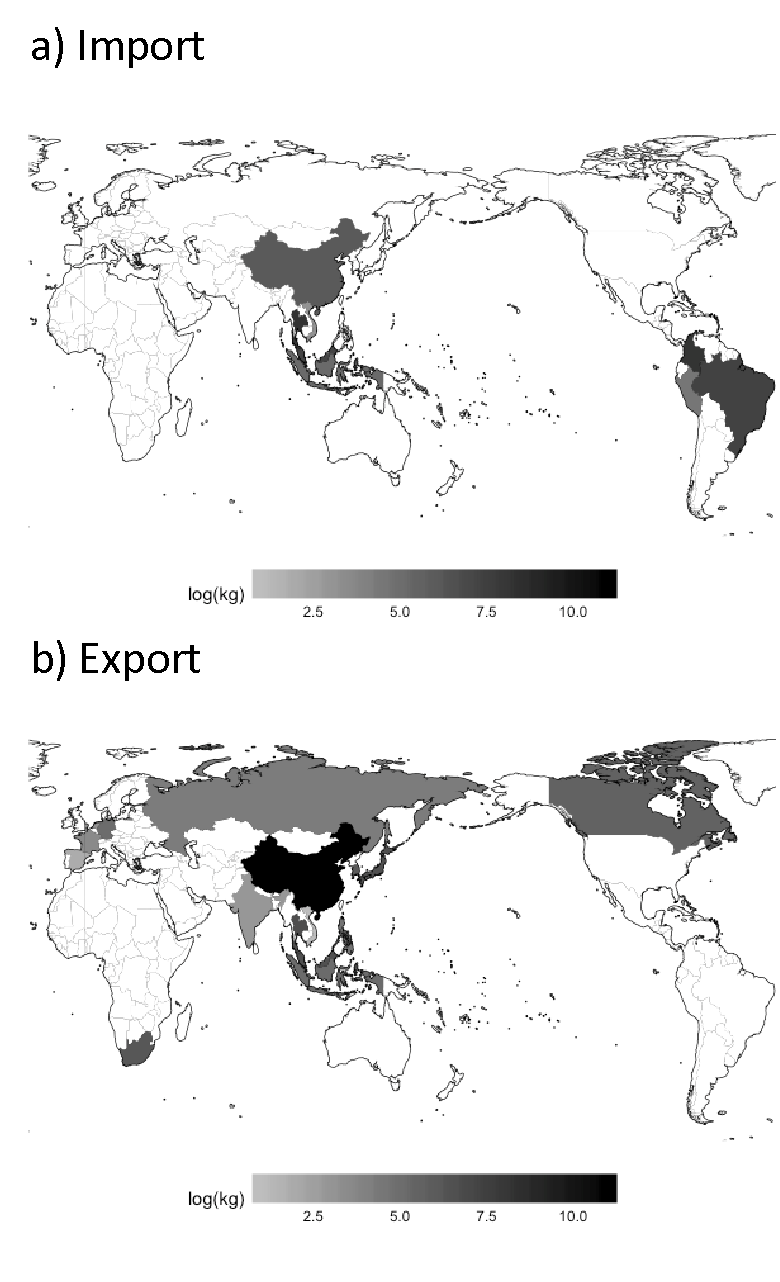

**Figure S4**. Map of countries with which Taiwan traded freshwater ornamental fish between 2013 and 2017. The grey-scaled bar on the bottom indicates the mean annual weight of fishes imported to Taiwan (a) and fishes exported from Taiwan (b).
